# From paralogy to hybridization: Investigating causes of underlying phylogenomic discordance using the complex genus *Packera* (Senecioneae; Asteraceae)

**DOI:** 10.1101/2023.08.14.553290

**Authors:** Erika R. Moore-Pollard, Jennifer R. Mandel

## Abstract

**Premise of the study:** Underlying discordance in phylogenomic studies is becoming more common, and the answer is not as simple as adding more data. Biological processes such as polyploidy, hybridization, and incomplete lineage sorting are main contributors to these issues and must be considered when generating phylogenies. Otherwise, interpretations of evolutionary relationships could be misleading.

**Methods:** To obtain a better understanding of potential gene flow and its effect on phylogenetic trees, we investigated the causes and consequences of nuclear discordance using the genus *Packera* to understand how they influence the phylogenetic patterns seen in this complex group. To do this, we compared the topology and support values of *Packera* phylogenies resulting from various paralog selection or pruning methods. We then investigated whether pruning the paralogs instead of performing a selection process affected the topology and support of our phylogeny. To investigate hybridization and its effect on species relationships in our tree, we used likelihood methods to infer phylogenetic networks to find any evidence of gene flow among species lineages in this complicated genus.

**Key results:** We found that performing different paralog selection or pruning methods does impact our understanding of the evolutionary relationships within *Packera*, and that addressing these paralogs with more rigorous methods than the typical pipeline increases concordance within the resulting phylogenies. Additionally, investigating reticulation events within highly discordant clades showed that ancestral hybridization and reticulation events are common throughout *Packera*.

**Conclusions:** Investigating underlying biological processes by testing various methods can provide further insight into complex species relationships and levels of discordance within phylogenomic studies.

## INTRODUCTION

The field of phylogenetics is drastically changing with the increased use of multi-locus and genomic data. A major challenge with this expansion is that conflicting genealogical histories often exist in different genes, and the answer is not as simple as increasing the amount of data (Degnan and Rosenberg, 2009). These differing gene histories can cause gene tree discordance that occurs when phylogenies obtained from individual gene trees differ among themselves and from the species tree (Mendes et al., 2016), potentially leading to incongruent tree topologies. Underlying discordance and uncertainty are frequently seen in phylogenomic studies and are repeatedly explained as results of biological processes including gene flow (hybridization and introgression), incomplete lineage sorting (ILS), or paralogous sequence duplication and loss (Pease et al., 2018).

Understanding gene flow between lineages is important in the evolutionary history of plants (Funk, 1985; Rose et al., 2021), especially when considering its role in speciation events. Speciation in plants mainly occurs through the process of hybridization, also known as “reticulate” speciation or reticulation (Coyne and Orr, 2004). Plants can speciate via hybridization in two ways: recombinational speciation, when a new, reproductively isolated species can be derived from two or more parental species, without a change in ploidy level, via hybridization (McCarthy et al., 1995), or by polyploid hybridization, when polyploids arise by interspecific hybridization or genome doubling (Ramsey and Schemske, 1998; Soltis et al., 2015). The latter, also known as allopolyploidy, is considered the most common and well known, and has received the greatest deal of attention.

Polyploidy, also known as whole-genome duplication (WGD), is considered an important evolutionary force and plays a major role in the evolution and speciation of many eukaryotic organisms. Polyploidy occurs in both animals (Evans et al., 2004; Gallardo et al., 2004; Braasch and Postlethwait, 2012; Nice et al., 2013) and plants (Vision et al., 2000; Wendel and Cronn, 2003; Peirson et al., 2012), with up to 70% of all angiosperms and 95% of monilophytes (ferns, horsetails, and club mosses; previously known as pteridophytes) having experienced polyploidization events (Grant, 1971; Masterson, 1994; Soltis and Soltis, 1999; Leebens-Mack et al., 2019). As a result, plant genomes frequently contain large numbers of paralogous sequences (hereafter referred to as “paralogs”) from repeated gene and genome duplication events (Lynch and Conery, 2000; Panchy et al., 2016; Morales-Briones et al., 2022).

During phylogeny reconstruction, the presence and abundance of paralogs presents a challenge when inferring orthology. Some ortholog recovery pipelines remove all genes that show evidence of potential paralogy (i.e., Phyluce; Faircloth, 2016). This approach may be appropriate if only a small number of genes show evidence of paralogy; however, in some datasets, pruning all paralogs could result in a significant reduction in the number of genes retained for phylogeny construction. Other phylogenomic pipelines do not remove paralogs but instead select the gene copy that is the longest or has the highest similarity to the target (i.e., HybPiper; Johnson et al., 2016), allowing the user to retain a larger number of genes for phylogenomic analyses. Given this, pipelines like these may be better suited for studying groups complicated by extensive polyploidy.

The evolution of plants is full of examples of polyploidy. In fact, it is hypothesized that all flowering plants (angiosperms) have experienced at least one polyploid event in their ancestry known as the zeta (ζ) event, which occurred prior to gymnosperms and angiosperms divergence about 340 MYA (Panchy et al., 2016; Alix et al., 2017). Additionally, plant groups across all taxonomic levels are predicted to have arisen via ancient hybridization events, such as the genus *Quercus* L. (McVay et al., 2017; Crowl et al., 2020), tribe Hippeastreae within Amaryllidaceae (García et al., 2014, 2017), and tribe Anthemideae within Compositae (Zhang et al., 2021), see Stull et al. (2023) for a full review.

Compositae, also known as Asteraceae, is known as the largest angiosperm family, making up 10% of all flowering plants (Palazzesi et al., 2022). Multiple WGD events have been suggested in Compositae (Barker et al., 2008, 2016; Huang et al., 2016; Zhang et al., 2020b), with the most recent number proposed being 41 (Zhang et al., 2021). These WGD events have occurred across all taxonomic levels of the family creating the immense diversity seen today. For example, previous studies have shown that the largest and most diverse tribe, Senecioneae, has an ancient WGD event that played a large role in its explosive diversification from South America to vast areas of the Earth via an African route (Barker et al., 2016; Huang et al., 2016; Mandel et al., 2019; Zhang et al., 2021; Palazzesi et al., 2022). Evolutionary relationships within Senecioneae have continuously changed (Pelser et al., 2007, 2010; Funk et al., 2009), leading researchers to hypothesize that reticulation and/or hybridization events have occurred among members of this tribe (Watson et al., 2020).

One genus within Senecioneae, *Packera* Á. Löve & D. Löve, is estimated to have about 88 species and varieties, and is known to be complicated by hybridization, polyploidy, and reticulation (Barkley, 1988; Trock, 2006; Moore-Pollard and Mandel, 2023). An estimated 40% of *Packera* taxa present polyploidy, aneuploidy, and other cytological disturbances (Barkley, 1988; Trock, 2006), complicating phylogenetic reconstruction of this group (Moore-Pollard and Mandel, 2023). Previous phylogenomic work by Moore-Pollard and Mandel (2023) showed high levels of underlying discordance within *Packera* given that only 49% of the gene trees were represented in the final species tree, and the remaining 51% of the gene trees show differing evolutionary relationships. Although not investigated, the authors predicted that extensive paralogy and hybridization are influencing the results and making resolving evolutionary relationships in the group complicated.

Thus, to obtain a better understanding of potential gene flow and its effect on complex systems such as *Packera*, we investigated the causes and consequences of nuclear discordance, including evolutionary processes of incomplete lineage sorting (ILS) and gene flow, to understand how they influence the phylogenetic patterns seen in *Packera*. To do this, we compared the topology and support values of *Packera* phylogenies resulting from the normal paralog selection processes defined by HybPiper, along with other paralog selection or pruning methods. We then investigated whether pruning the paralogs instead of performing a selection process affected the topology and support of our phylogeny. To investigate hybridization and its effect on the species relationships in our tree, we performed network analyses to discover any evidence for gene flow across species lineages in this complicated genus. In doing so, we aim to address: 1) does utilizing different paralog selection/pruning methods generate different results and provide higher resolution than typical methods, 2) is there evidence of ancient hybridization within *Packera*, and 3) can previous incongruence be explained by hybridization, introgression, ILS, or paralog duplication/loss? We anticipate that our approach of investigating the influence of paralogy and hybridization on underlying discordance in *Packera* will serve as a model in other complex plant groups complicated by hybridization and polyploidy.

## MATERIALS AND METHODS

### Specimen collection

A total of 108 taxa were used in this study, including 106 taxa previously sequenced and studied in Moore-Pollard and Mandel (2023). The remaining two taxa, *Packera paupercula* var. *paupercula* (Michx.) Á.Löve & D.Löve and *Packera subnuda* var. *subnuda* (DC.) Trock & T.M.Barkley, were added to this study and follow the DNA extraction and sequencing methods of Moore-Pollard and Mandel (2023). These two samples were then sequenced on an Illumina NovaSeq6000 at HudsonAlpha Institute of Technology (Huntsville, Alabama, USA). A complete list of sampled species, herbarium vouchers, and publication status can be found in Appendix S1 (see Supplemental Data with this article). Sequence stats were obtained using the *pxlssq* function in Phyx (Brown et al., 2017).

### Sequence processing and ortholog assembly

Raw FASTQ data were cleaned from adapter contamination and low-quality bases using the program Trimmomatic v. 0.36 (Bolger et al., 2014). Cleaned FASTQ data were then implemented in the HybPiper v. 2.0.1 (Johnson et al., 2016) pipeline to match the reads to the target loci contained in the Compositae-1061 probe set (Mandel et al., 2014). Paralog detection was carried out using the “paralog_retriever” command in HybPiper (https://github.com/mossmatters/HybPiper/wiki/Paralogs). This single command performs the function of previous scripts “paralog_investigator.py” and “paralog_retriever.py” from earlier versions of HybPiper, which flags loci that indicate potential paralogs, then extracts the assemblies that were flagged, respectively. Paired-end reads were then mapped to the target loci using BWA v. 0.7.17 (Li and Durbin, 2009), then assembled into contigs with specified kmer lengths (21, 33, 55, 77, 99) using SPAdes v. 3.5 (Bankevich et al., 2012). A multifasta was generated for each locus containing the sequences of that specific locus for all taxa in the run. These multifastas were aligned into “supercontigs” containing target and off-target sequence data, separately using MAFFT v. 7.407 (Katoh and Standley, 2013).

### Phylogenetic analyses

To compare the differences between paralog pruning and selection methods, we generated phylogenies utilizing three different pipelines that handle paralogs either using a selection or pruning process: ASTRAL-III (selection), ASTRAL-Pro (selection), and TEO (pruning; further details below; refer to Table 1 and Figure 1).

**Fig. 1.**
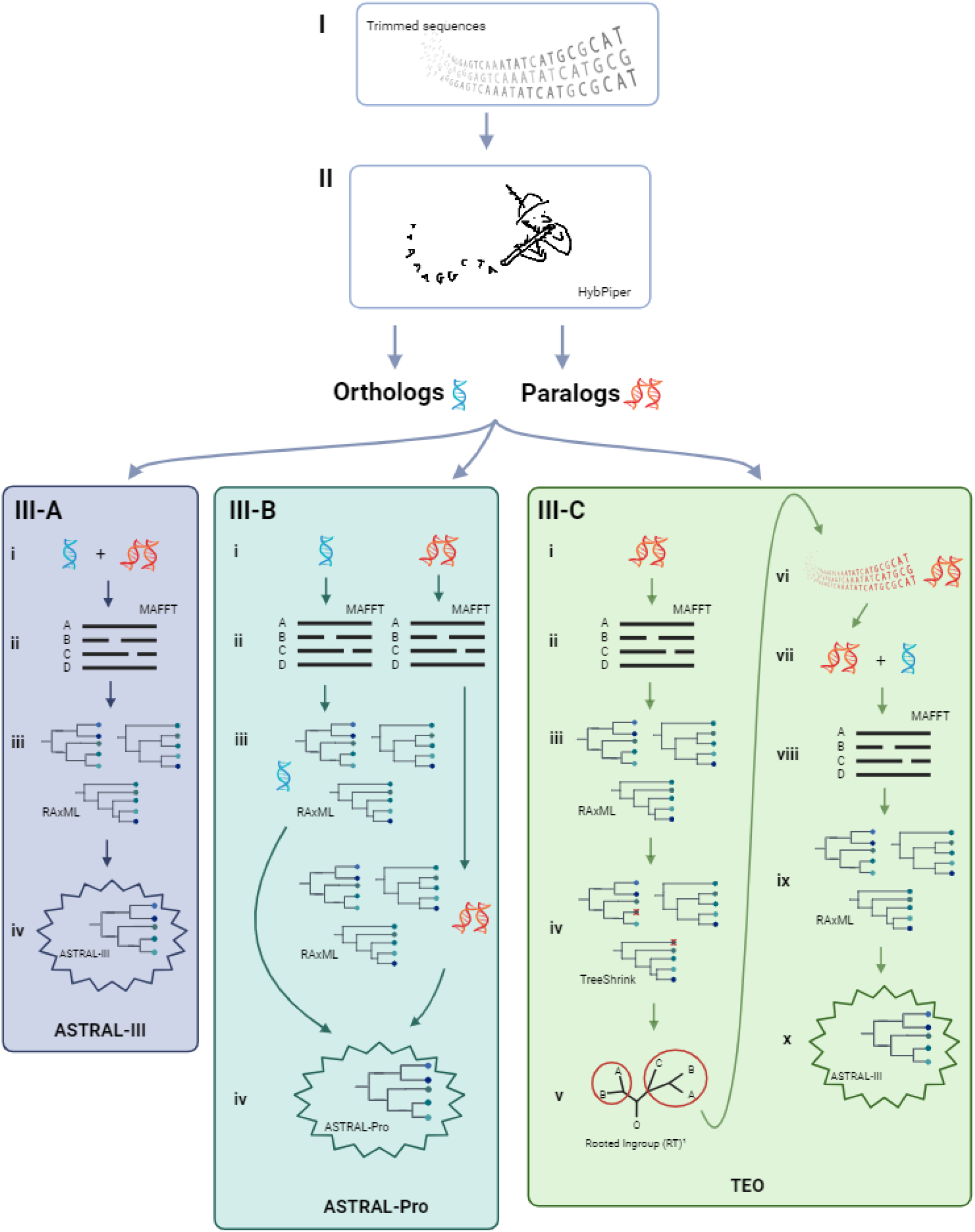
A visual of the pipelines used to perform paralog selection/pruning processes. First, raw sequences are cleaned and trimmed (**I**), then used as input to HybPiper to determine orthology (**II**). From there, the sequences go through the ASTRAL-III (**III-A**), ASTRAL-Pro (**III-B**), and TEO (**III-C**) pipelines. For ASTRAL-III (**III-A**): the orthologs and paralogs are combined (**i**) and aligned using MAFFT (**ii**) then used as input to RAxML to generate gene trees (**iii**). A final species tree is inferred using ASTRAL-III (**iv**). For ASTRAL-Pro (**III-B**): the orthologs and paralogs are separately (**i**) aligned using MAFFT (**ii**), then ran through RAxML to generate individual ortholog and paralog gene trees (**iii**), the resulting gene trees are then combined to generate the species tree using ASTRAL-Pro (**iv**). For TEO (**III-C**): only the paralogs (**i**) are aligned using MAFFT (**ii**) and gene trees are generated using RAxML (**iii**). The gene trees are then trimmed of spurious taxa using TreeShrink (**iv**). Resulting gene trees are then used as input to the rooted-ingroup (RT) method to prune paralogs (**v**). The resulting gene trees of RT are then converted to fasta sequences (**vi**) and combined with the original orthologs (**vii**) and aligned using MAFFT (**viii**), to make gene trees using RAxML (**ix**), and infer a final species tree using ASTRAL-III (**x**). Created with BioRender.com. ^1^ Yang and Smith (2015).

**Table 1.** Table detailing the differences between ASTRAL-III, ASTRAL-Pro, and TEO. Value explanations can be found in the text.

|  | **ASTRAL-III** | **ASTRAL-Pro** | **TEO** |
| --- | --- | --- | --- |
| Paralog modification | Selection | Selection | Pruning |
| # Taxa | 108 | 108 | 104 |
| # Paralogs | 810 | 810 | 810 |
| Modified Paralog Input # | 810 | 810 | 782 |
| Modified Paralog Final # | 809 | 809 | 692 |
| Final Locus # | 1,049 | 1,049 | 932 |
| Species Tree Inference Method | ASTRAL-III | ASTRAL-Pro 2 | ASTRAL-III |
| Normalized Quartet Score | 0.489 | N/A | 0.482 |
| # Concordant Nodes | 9 | 9 | 5 |
| # Discordant Nodes | 25 | 28 | 29 |

*ASTRAL-III —* The first pipeline, hereafter referred to as ASTRAL-III, followed a typical pseudo-coalescent approach where no paralogs were pruned, but paralog selection followed the default settings in HybPiper. HybPiper’s paralog selection process chooses the contig with the greatest coverage depth. If the contigs have similar depth, then the sequence with the greatest percent identity to the reference is chosen (Johnson et al., 2016). For this analysis, the aligned supercontigs were used to generate single-copy gene trees with RAxML v. 8.1.3 (Stamatakis, 2014) with 1,000 bootstrap (BS) replicates under the GTR+I+Γ nucleotide substitution model. These gene trees were used as input to ASTRAL-III v. 5.7.3 (Zhang et al., 2018) to generate a species tree with local posterior probability (LPP) values. To determine the proportion of gene trees found in the final species tree, the normalized quartet score was calculated using the –q option in ASTRAL-III. Additionally, the –k option was used to obtain branch lengths.

*ASTRAL-Pro —* Another paralog selection method implemented in this study utilized ASTRAL-Pro2 (ASTRAL for PaRalogs and Orthologs) v1.13.1.3 (Zhang et al., 2020a; Zhang and Mirarab, 2022), hereafter referred to as ASTRAL-Pro. This tool provides a quartet-based species-tree inference method, and unlike ASTRAL-III, ASTRAL-Pro allows multi-copy gene trees to be used as input, further accounting for orthology and paralogy. In this case, paralog selection allows all paralogous sequences to be used as input to generate the final species tree instead of selecting a single paralog to represent an individual gene’s history.

The process to generate a final species tree for the ASTRAL-Pro pipeline is similar to ASTRAL-III as described above but differs by having the user generate separate alignment and maximum likelihood analyses on the genes that are flagged as paralogs during the HybPiper run and on the single-copy contigs, separately. Single-copy and paralogous gene trees were generated under the GTR+I+Γ nucleotide substitution model in RAxML with 1,000 BS replicates. The RAxML output files were then used as input for ASTRAL-Pro2 to generate the final species tree with LPP values at each node.

*TEO —* Instead of selecting paralogs, the third approach prunes paralogs by utilizing the Target Enrichment Orthology (TEO) pipeline (https://bitbucket.org/dfmoralesb/target_enrichment_orthology/src/master/), which investigates the flagged loci from HybPiper and determines which loci can be retained or pruned given specified conditions (Yang and Smith, 2014; Morales-Briones et al., 2022). Only loci flagged as paralogous from HybPiper were used as input for the TEO pipeline. Briefly, individual paralogous sequences were aligned using MAFFT with default settings. Aligned columns containing more than 90% missing data were removed using Phyx. Individual gene trees were generated using RaxML with 100 BS replicates under the GTR+I+Γ nucleotide substitution model. Only tips with the highest number of unambiguous characters were retained for each gene, while mono– and paraphyletic tips belonging to the same taxon were masked. Rogue tips were removed with TreeShrink v. 1.3.2 (Mai and Mirarab, 2018) using default settings.

Ortholog inference was then carried out on the resulting sequences using an outgroup-aware strategy originally proposed by Yang & Smith (2014) coined the “rooted ingroup” (RT) approach. This approach searches for subtrees with at least 25 ingroup taxa and roots them by the designated outgroups (here Tribe Anthemideae: *Achillea nobilis* L., *Anthemis arvensis* L., *Artemisia vulgaris* L., and *Cotula turbinata* L.), then removes the ingroup taxa as rooted trees. Inferred gene duplications, or paralogs, are then removed from root to tip in the rooted tree. This RT method maximizes the number of orthologs retained since it allows for non-monophyletic and duplicated taxa in the outgroups (Yang and Smith, 2014; Morales-Briones et al., 2022). Other orthology inference methods proposed by Yang & Smith (2014), the “maximum inclusion” (MI) and “monophyletic outgroup” (MO) methods, could not be used in our study since our outgroup taxa are known to have whole genome duplications (Zhang et al., 2021; Palazzesi et al., 2022).

The resulting ortholog trees from TEO, along with the remaining loci that were not originally flagged as paralogous, were then converted to fasta files, and used as input for generating a final phylogenetic tree. Since the TEO orthologs did not contain the four Anthemideae outgroup species, those taxa were pruned from the originally orthologous loci from HybPiper using the *pxrmt* function in Phyx, totaling 104 taxa in the final TEO tree. Species tree inference then followed ASTRAL-III’s protocol described above.

All resulting trees from ASTRAL-III, ASTRAL-Pro, and TEO were visualized using the package *phytools* (Revell, 2012) in R v. 4.0.5 (R Core Team, 2016; RStudio, 2020).

### Measuring discordance

All final species trees generated using ASTRAL-III, ASTRAL-Pro, and TEO were analyzed for underlying discordance using Quartet Sampling (Pease et al., 2018), which assesses the confidence, consistency, and informativeness of internal tree relationships, and the reliability of each terminal branch. Quartet Sampling evaluates if a phylogeny has a lack of support due to lack of information, discordance from ILS or introgression, or misplaced or erroneous taxa. Three scores are calculated (QC, QD, QI) for each internal branch of the focal tree which together allows different interpretations of the branches, as well as a Quartet Fidelity (QF) score that reports how frequently a taxon is included in concordant topologies. Quartet Sampling efficiently synthesizes phylogenetic tests and offers more comprehensive and specific information on branch support than conventional measures (i.e., bootstrap or local posterior probability). Quartet Sampling requires a final rooted tree and a concatenated matrix of all aligned gene trees. The final ASTRAL-III and ASTRAL-Pro trees were rooted to the four outgroup Anthemideae taxa (*Achillea nobilis*, *Anthemis arvensis*, *Artemisia vulgaris*, and *Cotula turbinata*), while the TEO tree was rooted to *Doronicum pardalianches* L. using the *pxrr* function in Phyx. The final concatenated gene matrices were generated using FASconCAT-G v. 1.02 (Kück and Longo, 2014).

Additionally, *bellerophon* (https://git.sr.ht/~hms/bellerophon) was used to quantify and visualize discordance in the three final species trees. The program *bellerophon* is similar to PhyParts (Smith et al., 2015), but can more easily handle large datasets. Final images containing pie charts indicating whether gene trees were conflicted, supported, or uninformative at each node were generated using the program *gokstad* (https://git.sr.ht/~hms/gokstad). The program *bellerophon* requires using rooted gene trees and a rooted, final species tree as input. Rooting gene trees for *bellerophon* decreased the number of genes used as input in each analysis: ASTRAL-III and ASTRAL-Pro had 142 and 325 genes (out of 1,049), respectively, that did not contain at least one of the outgroup Anthemideae taxa and were deleted. The TEO analysis had 451 genes (out of 932) that did not contain the outgroup taxon *Doronicum pardalianches* and were removed from the analysis. Resulting in 907 (ASTRAL-III), 724 (ASTRAL-Pro), and 481 (TEO) gene trees used as input for *bellerophon*.

### Tree topology comparison

To evaluate similarity among the ASTRAL-III, ASTRAL-Pro, and TEO tree topologies, we calculated the adjusted Robinson-Foulds (RF_adj_) distance as outlined by Mitchell et al. (2017) between the two trees using the RF.dist() function in package *phangorn* (Schliep, 2011) in R v. 4.0.5 (R Core Team, 2016; RStudio, 2020), using unrooted trees as input. The Anthemideae outgroup taxa were trimmed from the trees in R using the drop.tip() function in package *ape* (Paradis and Schliep, 2019) for equal comparisons between ASTRAL-III, ASTRAL-Pro, and TEO. The RF distance was adjusted by setting the “normalize” argument to TRUE, which divides RF by the maximal possible distance as described in Steel and Penny (1993). Given that the input trees are unrooted and binary, this value is then 2*n*-6, where *n* is the number of nodes on the tree (resulting equation: RF_adj_ = RF/(2*n*-6)). The RF_adj_ values range from zero to one, indicating whether the tree topologies are identical or completely dissimilar, respectively.

### Reticulation

The Species Networks applying Quartets (SNaQ) function in the Julia v. 1.8.1 (Bezanson et al., 2017) package PhyloNetworks v. 0.15.2 (Solís-Lemus and Ané, 2016; Solís-Lemus et al., 2017) was used to investigate any evidence for reticulate evolution among *Packera* taxa. This package uses unrooted, gene trees to infer phylogenetic networks in a pseudolikelihood framework with multi-locus data, while accounting for ILS and reticulation. We tested the fit of multiple models and gradually increased the number of reticulation events (h) from 0 to 5 to find which pseudolikelihood score reached a plateau, given recommendations by Solís-Lemus et al. (2017). This process is computationally intense, so only clades containing 14 taxa or less that had three or more discordant nodes in the ASTRAL-III tree, as determined by a negative Quartet Concordance (QC) score resulting from Quartet Sampling, were used as input for PhyloNetworks. The remaining taxa were pruned from the tree using the *pxrmt* function in Phyx. Hybridization networks were visualized using Dendroscope v.3.7.6 (Huson and Scornavacca, 2012).

## RESULTS

Illumina sequencing resulted in a total of 2.9 billion reads and 19 million sequences across the two newly sequenced taxa in this study. *Packera paupercula* var. *paupercula* averaged around 200 million reads and 1.3 million sequences, and *P. subnuda* var. *subnuda* resulted in 2.7 billion reads and 18 million sequences. Raw data for *P. paupercula* var. *paupercula* and *P. subnuda* var. *subnuda* are deposited in NCBI as SRR24864449 and SRR24895390, respectively. HybPiper recovered 1,057 out of the 1,061 targeted loci on the 108 samples and flagged 810 loci for paralogs. Summary statistics can be found in Appendix S2.

### Summary statistics

All 1,057 loci were used as input for the ASTRAL-III and ASTRAL-Pro trees. For ASTRAL-III, eight loci were removed from further analyses because the resulting RAxML tree was empty or contained three or fewer taxa, resulting in 1,049 total loci used as input to generate the final ASTRAL-III species tree. For ASTRAL-Pro, MAFFT and RAxML were run on the 810 paralog and 251 ortholog fasta files independently. Similar to the ASTRAL-III tree, loci that were missing or had three or fewer taxa were removed from further analyses, resulting in 809 paralog and 240 ortholog loci used as input for the RAxML run. The RAxML output files were then combined, totaling 1,049 gene files, and used as input for ASTRAL-Pro to generate a final species tree.

All 810 paralogous loci flagged by HybPiper were used as input for the TEO pipeline. Aligning and trimming paralogous loci prior to generating homolog trees removed 28 loci, resulting in 782 loci being used as input for the RAxML run. Additionally, three genes had taxa containing spurious tips that were removed by TreeShrink. The end of the RT pipeline resulted in 692 final fastas generated from ortholog trees. The remaining 118 loci originally flagged for paralogs were then pruned from further analyses. The 240 loci that originally did not have paralog warnings were added to the final RT fastas, but had the four, outgroup Anthemideae taxa removed from all gene trees, totaling 932 loci and 104 taxa used as input to generate a final species tree with RAxML and ASTRAL-III. A locus list with their ortholog/paralog status can be found in Appendix S3.

A summary of the three final species trees can be found in Table 1.

### Topological comparisons and discordance

The 108 taxa ASTRAL-III tree is slightly incongruent to the original tree containing 106 taxa published by Moore-Pollard and Mandel (2023) (RF_adj_ = 0.17; Appendix S4). Additionally, the number of gene trees represented in the final species tree containing 108 taxa is slightly lower than the 106 taxa tree at 0.489 compared to 0.493 from the previous study (Moore-Pollard and Mandel, 2023), though both indicate that about 49% of the gene trees match the final species tree. Support values varied depending on which paralog process was used (Figs. 2 – 4). Phylogenies containing all LPP values can be found in Appendices S5 – S7.

**Fig. 2.**
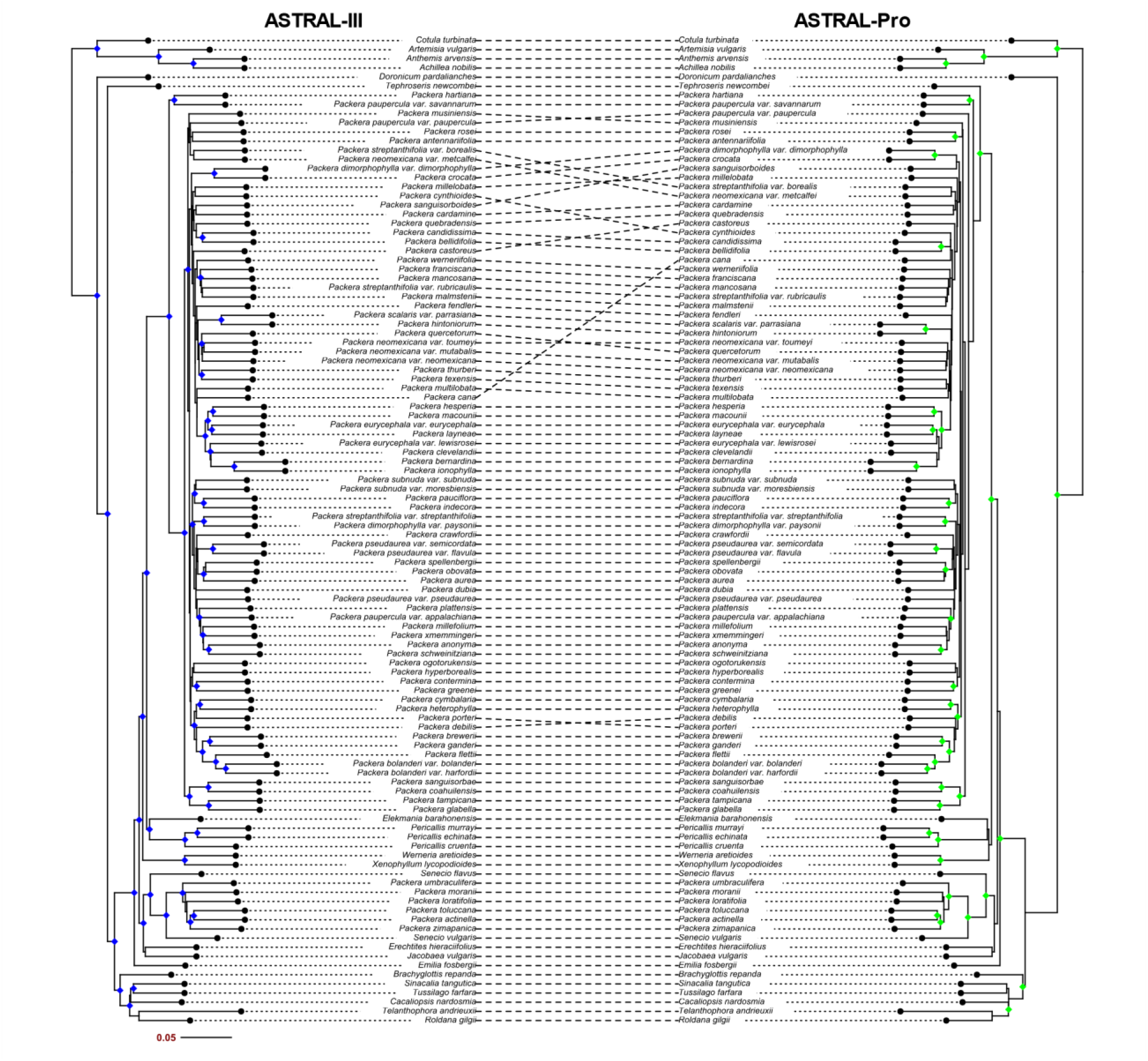
Tanglegram of ASTRAL-III (left) and ASTRAL-Pro tree (right) of all 108 taxa. Nodes that are considered highly supported (LPP > 0.90) are indicated by a blue diamond (ASTRAL-III) or green diamond (ASTRAL-Pro).

The ASTRAL-Pro and TEO tree topologies were most similar (RF_adj_ = 0.347), with the ASTRAL-III tree as equally dissimilar to ASTRAL-Pro and TEO trees (RF_adj_ = 0. 366) (Table 2, Figs. 2 – 4). Quartet Sampling indicated that discordance remains high independent of which method was used to handle paralogy (selection or pruning). ASTRAL-III and ASTRAL-Pro had the most concordant and highly supported nodes (QS = 1/NA/1) when compared to TEO, and ASTRAL-III had the overall lowest number of discordant nodes (QC < 0; Table 1). However, ASTRAL-Pro had the most concordant number of nodes within our focal group, *Packera*. Thus, if nodes from the outgroup Anthemideae taxa were not considered, ASTRAL-Pro had the highest number of concordant nodes compared to ASTRAL-III and TEO (Fig. 5). See Appendices S8 – S10 for all Quartet Sampling results.

**Table 2.** Pairwise adjusted Robinson-Foulds (RF_adj_) distance values between the three different tree topologies: ASTRAL-III, ASTRAL-Pro, and TEO. Values closer to zero indicate that the tree topologies are completely identical, values closer to one indicate completely dissimilarity.

|  | ASTRAL-III | ASTRAL-Pro | TEO |
| --- | --- | --- | --- |
| ASTRAL-III | 0 | --- | --- |
| ASTRAL-Pro | 0.3663366 | 0 | --- |
| TEO | 0.3663366 | 0.3465347 | 0 |

A similar interpretation can be made with the *bellerophon* results. The resulting pie charts indicate that the ASTRAL-Pro and TEO trees have fewer conflicting gene trees than ASTRAL-III indicated by the red portions of the pie charts at each node and the tip labels being red/orange. Alternatively, ASTRAL-Pro and TEO seem to have more uninformative gene trees than ASTRAL-III given the higher number of pie charts showing green portions and tip labels being yellow/green (Fig. 6). It is important to note that these uninformative branches could be influenced by support in the underlying gene trees, such as the second-best alternative topology having strong support.

### Reticulation events

Four *Packera* clades containing 14 or less taxa were determined to be highly discordant (three or more nodes had negative QC values) given results of Quartet Sampling on the ASTRAL-III tree (Fig. 5) and were investigated for hybridization or reticulation events using PhyloNetworks. Additionally, a single species from each major clade was then used to represent backbone relationships throughout the entire phylogeny, totaling five separate PhyloNetwork runs that will be referenced as: Backbone: representing 11 taxa that span the entire phylogeny; Eastern: a clade of 14 species either distributed along eastern North America or have known affiliations with those species; California/Mexico: ten taxa with distributions within California or Mexico; Rocky: a clade of five taxa that are distributed in the Central US, most within the Rocky Mountains; and Arctic/Alpine: 13 taxa that are present in colder climates along the Pacific Northwest and into Canada (Appendix S11). Log likelihood values indicated that the most likely number of reticulation events (h) varied depending on the run, with Backbone, California/Mexico, and Eastern showing the highest number of reticulation events (h = 5) and Rocky showing the lowest (h = 1). Arctic/Alpine was the second lowest with h = 2 (Appendix S11).

## DISCUSSION

In this study, we investigated how paralogy impacts phylogenetic reconstruction and interpretation within the genus *Packera*. We found that performing different paralog selection or pruning methods to determine orthology (ASTRAL-III, ASTRAL-Pro, and TEO) impacts our understanding of the evolutionary relationships within *Packera*, and that addressing these paralogs with more rigorous methods than the typical pipeline (ASTRAL-III) increases concordance across the resulting phylogenies. We also found that addressing paralogy did not fully resolve species relationships, potentially from the influence of reticulate evolution as many taxa were found to be influenced by hybridization or reticulation events.

### Discordance

Discordance varied depending on which paralog selection/pruning method was used. Overall, ASTRAL-III was the most concordant and highly supported, with ASTRAL-Pro as the next most supported. ASTRAL-Pro had the most concordant nodes within our focal group, *Packera*, compared to ASTRAL-III or TEO (Fig. 5), with TEO performing the worst out of all tested scenarios. One reason TEO may have underperformed is from the differences in locus retention between ASTRAL-III and ASTRAL-Pro methods when compared to TEO (1,049 vs. 932 loci retained, respectively; Table 1). Additionally, the RT method utilized in the TEO pipeline naturally loses information since it removes outgroups to determine orthology (Morales-Briones et al., 2022), providing another potential explanation for the differences between paralog selection and pruning methods. Even so, discordance remained high in all trees (Figs. 5 & 6), indicating that some other biological processes, such as hybridizations or reticulations, may be impacting our results.

### Tree topologies

Although ASTRAL-III and ASTRAL-Pro had more similar discordance values, ASTRAL-Pro and TEO shared the most similar topologies when compared to ASTRAL-III overall (Table 2, Figs. 2 – 4), demonstrating that ASTRAL-Pro and TEO present more similar topology results than ASTRAL-III; however, the difference is minor (Table 2). Additionally, the topologies of ASTRAL-Pro and TEO are more similar to the 106 taxa tree from Moore-Pollard and Mandel (2023). Given the tree topology and discordance results, ASTRAL-Pro appears to be the best fitting method for *Packera*. Even so, this trend may not be consistent with other complex plant groups, so we suggest testing multiple paralog selection/pruning methods on one’s dataset to determine the impact of paralogs on your tree topologies and support values.

Interestingly, most of the species associated with conflicting tree topologies are found in clades that were not considered highly supported, though not discordant. For example, the ASTRAL-III tree has a clade with low-support composed of *Packera musiniensis* (S.L.Welsh) Trock, *P. streptanthifolia* var. *borealis* (Torr. & A.Gray) Trock, *P. neomexicana* var. *metcalfei (Greene)* Trock & T.M.Barkley., *P. paupercula* var. *paupercula*, *P. rosei* (Greenm.) W.A.Weber & Á.Löve, and *P. antennariifolia* (Britton) W.A.Weber & Á.Löve. These same species show substantially different relationships in the TEO tree (Fig. 3). Even so, only one node in that ASTRAL-III clade was considered discordant (Fig. 5A). Similarly, *P. rosei* and *P. antennariifolia*, two species previously predicted to be influenced by long-branch attraction (Moore-Pollard and Mandel, 2023), demonstrate considerably different relationships within the trees dependent on sampling and paralog selection/pruning method (Figs. 3 & 4).

**Fig. 3.**
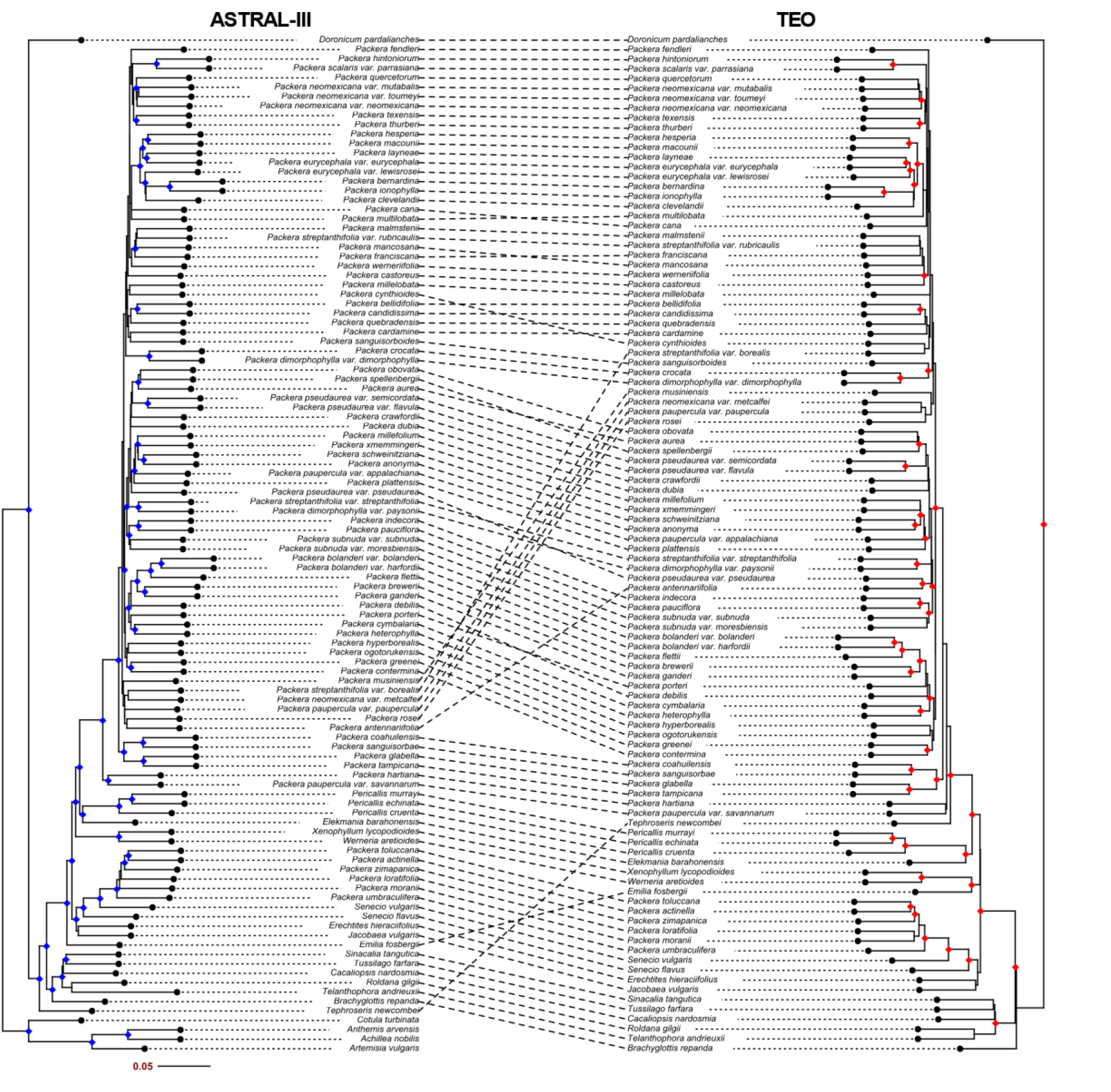
Tanglegram of ASTRAL-III (left) and ASTRAL-III tree of loci generated from TEO pipeline (right) of all 108 taxa. Nodes that are considered highly supported (LPP > 0.90) are indicated by a blue diamond (ASTRAL-III) or red diamond (TEO).

**Fig. 4.**
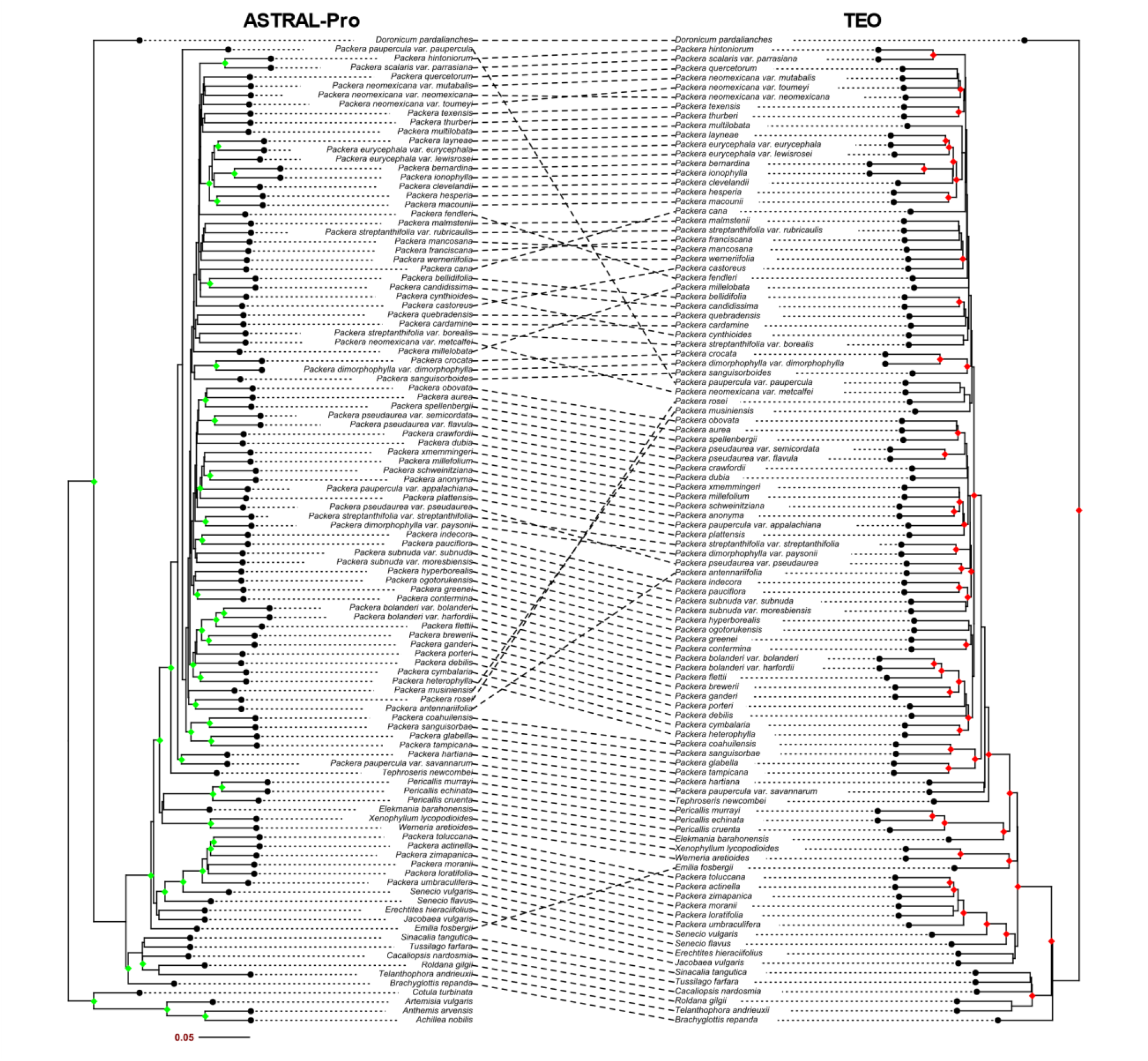
Tanglegram of ASTRAL-Pro (left) and ASTRAL-III tree of loci generated from TEO pipeline (right) of all 108 taxa. Nodes that are considered highly supported (LPP > 0.90) are indicated by a green diamond (ASTRAL-Pro) or red diamond (TEO).

We note that some *Packera* species are recovered in the outgroup instead of with the ingroup *Packera* species, particularly *P. actinella* (Greene) W.A.Weber & Á.Löve, *P. loratifolia* (Greenm.) W.A.Weber & Á.Löve, *P. toluccana* (DC.) W.A.Weber & Á.Löve, *P. umbraculifera* (S.Watson) W.A.Weber & Á.Löve, *P. zimapanica* (Hemsl.) C.C.Freeman & T.M.Barkley, and *P. moranii* (T.M.Barkley) C.Jeffrey. These species are thought to be misidentified as *Packera* species and have not yet been formally classified out of *Packera* (Barkley, 1985; Bain and Jansen, 1995; Bain et al., 1997; Bain and Golden, 2000; Pelser et al., 2007). A full description of these relationships can be found in Moore-Pollard and Mandel (2023).

Performing different paralog selection methods or pruning paralogs changed the sister relationship to *Packera*. For example, ASTRAL-Pro and TEO resulted in *Tephroseris newcombei* (Greene) B.Nord. & Pelser being sister to *Packera* with high support (0.99LPP), while ASTRAL-III has *Pericallis* D.Don and *Elekmania* B.Nord. as most closely related to *Packera* with high support (0.98LPP; Fig. 4). *Tephroseris newcombei* was once placed into *Packera* as *P. newcombei* (Greene) W.A.Weber & Á.Löve (Weber and Löve, 1981), but there is a general consensus that it should instead be placed in the genus *Tephroseris* (Rchb.) Rchb. (Janovec and Barkley, 1996; Bain and Golden, 2000; Golden et al., 2001; Nordenstam and Pelser, 2011). Interestingly, Moore-Pollard and Mandel (2023) found that *T. newcombei* placed within or sister to *Packera* with the plastid data but placed outside of Senecioneae with the nuclear data, similar to the ASTRAL-Pro and TEO results in this study. *Tephroseris* is taxonomically complicated as the placement of it within Senecioneae has continuously changed (Nordenstam and Pelser, 2011), so much so that it is sometimes placed within a separate subtribe, Tephroseridinae, along with two other genera, *Sinosenecio* B. Nord. and *Nemosenecio* (Kitam.) B. Nord. (Jeffrey and Chen, 1984). Even so, it is noteworthy that changing the paralog selection/pruning methods and/or data type (nuclear vs. plastid) drastically changes the relationship between *Packera* and *Tephroseris*.

### Reticulation events

Investigating hybridization events within highly discordant clades showed that ancestral hybridization and reticulation events may be common throughout *Packera*, as every tested clade had at least one event. This is unsurprising as botanists have previously predicted the presence of ancient hybridization, reticulation, and ILS events within this group (Bremer, 1994; Bain et al., 1997; Golden and Bain, 2000). The ancient hybridization events found can potentially explain the higher discordance seen within these clades, as hybridization, among other biological processes, are commonly associated with gene tree discordance (Hahn, 2007; Degnan and Rosenberg, 2009; Mendes et al., 2016; Morales-Briones et al., 2021; McLay et al., 2023). The results from the hybridization network analyses of the Backbone and four most discordant clades (Eastern, California/Mexico, Arctic/Alpine, and Rocky; Fig. 7) are discussed below.

**Fig. 5.**
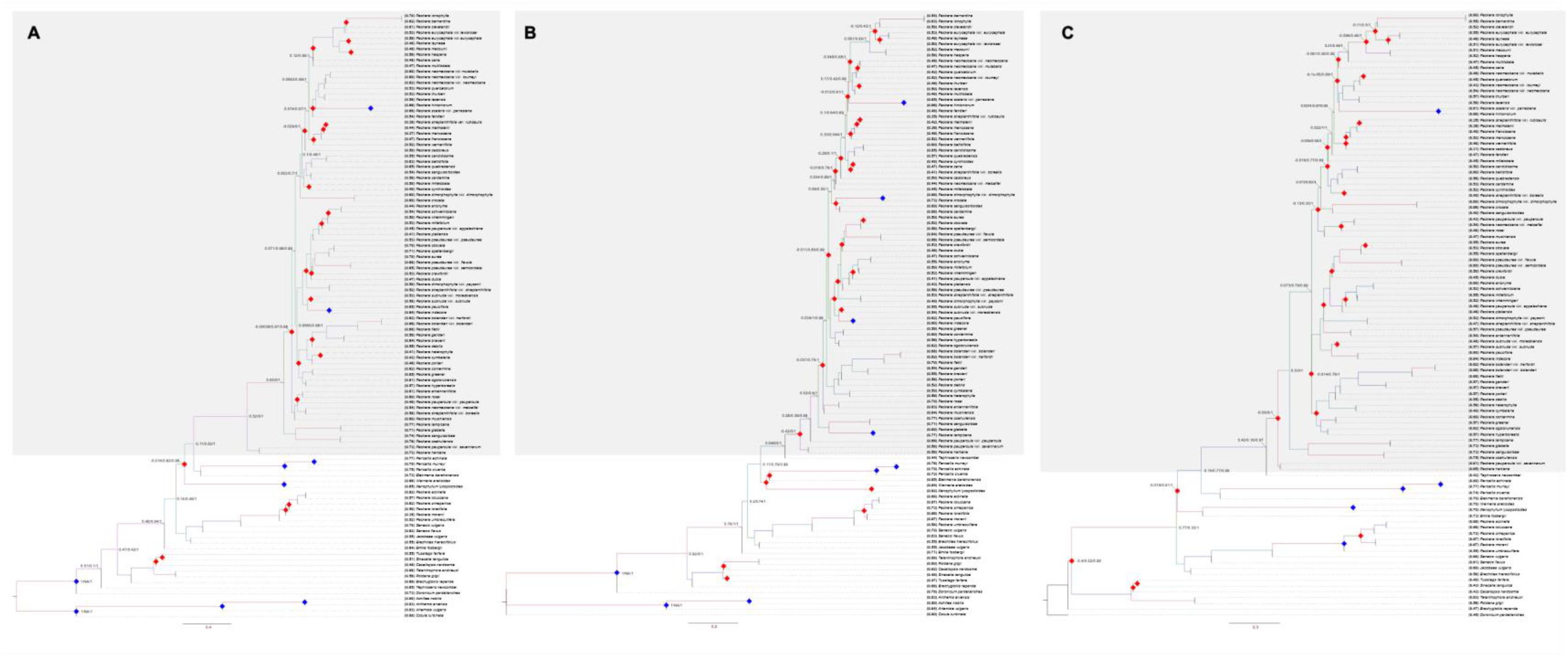
Results of Quartet Sampling on the (A) ASTRAL-III, (B) ASTRAL-Pro, and (C) TEO trees. *Packera* taxa are shaded gray. Nodes considered fully supported and concordant (1/NA/1) are indicated with a blue diamond. Nodes considered weakly supported and discordant (QC < 0) is indicated with a red diamond. QC/QD/QI values along the backbone of the phylogeny remain.

**Fig. 6.**
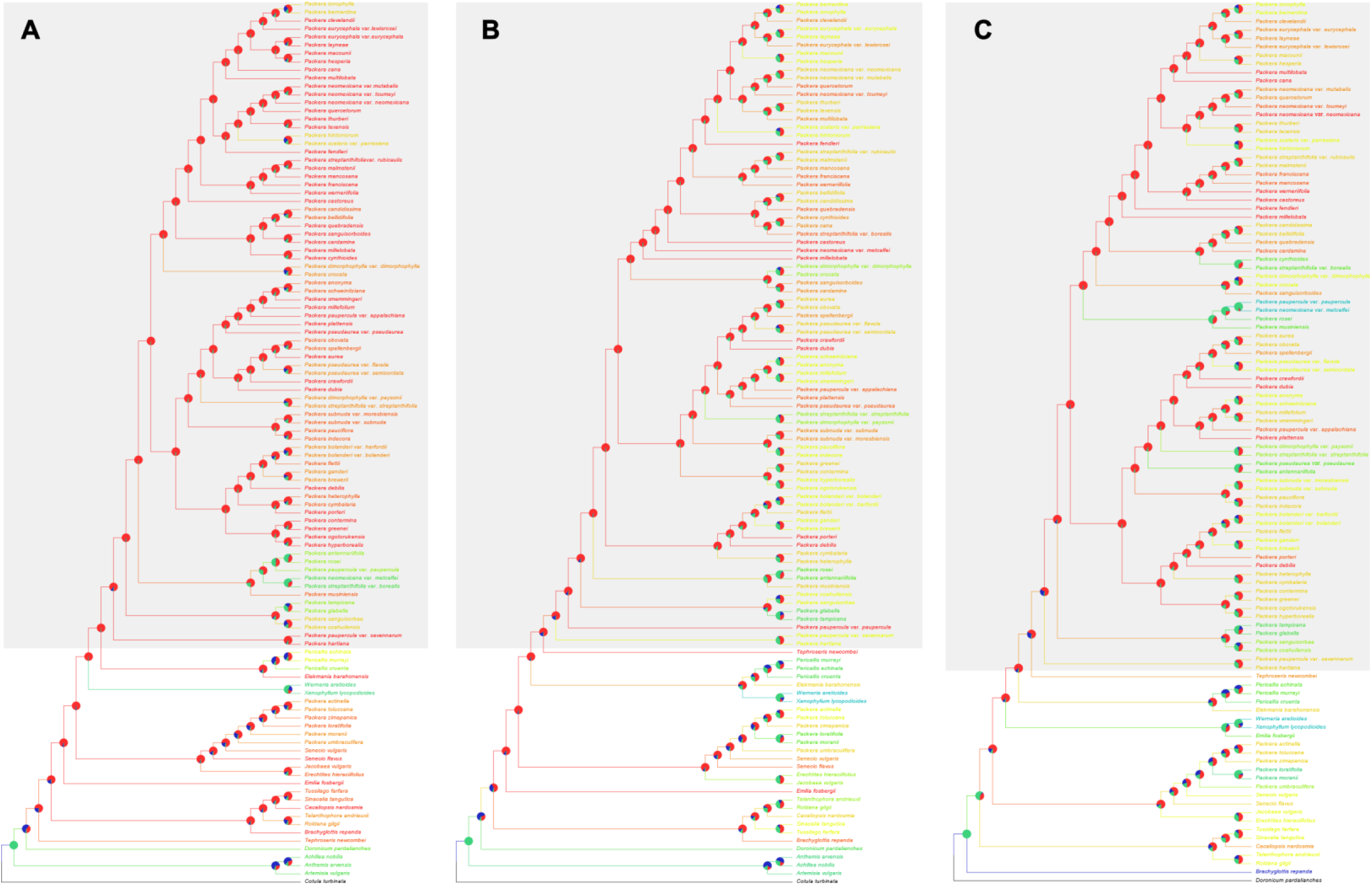
Results of *bellerophon* on the (A) ASTRAL-III, (B) ASTRAL-Pro, and (C) TEO trees. *Packera* taxa are shaded gray. Pie charts at each node indicate the proportion of gene trees that are supported (blue), conflicting (red) or lack information (green). Branches and tip labels are colored as a gradient to the proportion of conflicted gene trees: red/orange indicate a larger proportion of conflicting gene trees, yellow/green indicate a larger proportion of uninformative branches, while blues indicate more support. Black tip labels and branch lengths show the rooted taxon.

**Fig. 7.**
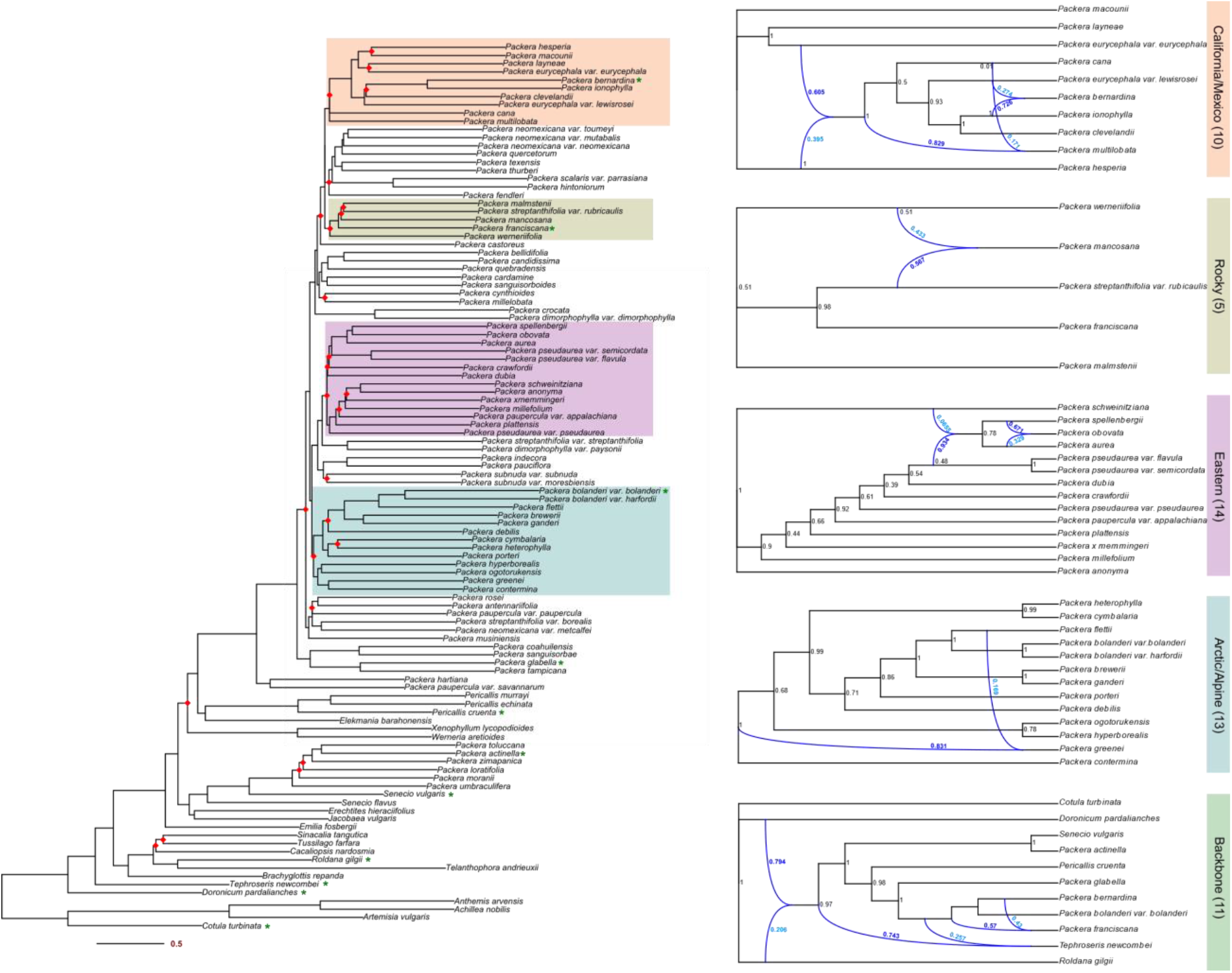
ASTRAL-III phylogeny of 108 taxa. Red diamonds at the node indicate discordance (QC < 0) given results from Quartet Sampling. Clades are highlighted according to majorly discordant clades and follow the coloring scheme of the PhyloNetwork networks to the right. Green stars to the right of some taxon names in the ASTRAL-III phylogeny relate to species included in the Backbone PhyloNetwork analysis. Black numbers in the PhyloNetwork networks indicate bootstrap support values, while light blue and dark blue numbering shows values of the hybrid edges.

#### Backbone

The Backbone network analysis shows a high proportion of reticulation events within the designated clade. Given that the number of species sampled in this run is low and covers a narrow range of the entire phylogeny, we do not suggest deeply interpreting the results. Instead, we want to point out what we believe is the biggest takeaway from this analysis, which is the ancestral reticulation event between the lineages that gave rise to *Roldana gilgii* (Greenm.) H.Rob. & Brettell and *Doronicum pardalianches* L., ultimately producing the remaining Senecioninae taxa (Fig. 7). This is particularly fascinating because researchers have previously predicted that *Packera*, among other Senecioninae taxa, resulted from a “Tussilaginoid” influence in its history given that they possess helianthoid type pollen, the most common pollen type in subtribe Tussilagininae s.str., instead of senecioid pollen which is typical of most Senecioneae members (Bain et al., 1997; Bain and Golden, 2000). Therefore, these results may provide additional support for this hypothesis that *Packera*, and additional Senecioneae members, formed via a “Tussilaginoid” influence.

#### Eastern

Hybridization results in the Eastern clade indicate a potential ancestral reticulation event between the lineages that gave rise to *Packera schweinitziana* (Nutt.) W.A.Weber & Á.Löve and *P. pseudaurea* (Rydb.) W.A.Weber & Á.Löve varieties, leading to the ancestors of *P. spellenbergii* (T.M.Barkley) C.Jeffrey, *P. obovata* (Willd.) W.A.Weber & Á.Löve, and *P. aurea* (L.) Á.Löve & D.Löve. Within that reticulation event is a nested hybridization event between *P. spellenbergii* and *P. aurea*, leading to the potential hybrid origin of *P. obovata* (Fig. 7). These results are interesting because *P. spellenbergii*’s placement in this Eastern clade has been questioned since *P. spellenbergii* is a species endemic to the Western United States in only three counties in New Mexico and Utah; quite geographically distinct from the remaining members in this clade. Therefore, having a connection from a potential ancestral hybridization event could provide an explanation for the species’ disjunct placement.

Curiously, an accepted hybrid species, *Packera* × *memmingeri* (Britton ex Small) Weakley, does not appear to have any evidence of an ancestral hybrid origin between its proposed parents, the widespread *P. anonyma* (Alph.Wood) W.A.Weber & Á.Löve and endangered *P. millefolium* (Torr. & A.Gray) W.A.Weber & Á.Löve (Uttal, 1984; Gramling, 2006; Weakley et al., 2011). This may be because its formation is more recent and cannot be distinguished with any phylogenetic signal, indicating that future work should investigate more recent hybridization events between species.

#### California/Mexico

The California/Mexico clade has the highest amount of predicted reticulation events, next to the Backbone clade. First, there is an ancestral reticulation event between the lineages that gave rise to *Packera hesperia* (Greene) W.A.Weber & Á.Löve and *P. eurycephala* var. *eurycephala* (Torr. & A.Gray) W.A.Weber & Á.Löve, leading to the formation of a small clade composed of mainly California endemic species or species with distributions throughout California. There is another anticipated ancestral reticulation event between the lineages that gave rise to *P. eurycephala* var*. lewisrosei* (J.T.Howell) J.F.Bain and *P. ionophylla* (Greene) W.A.Weber & Á.Löve, forming the rare and threatened *P. bernardina* (Greene) W.A.Weber & Á.Löve (Fig. 7). *Packera bernardina* and *P. ionophylla* are found in the same mountain range (San Bernardino Mountains, CA), while *P. eurycephala* var. *lewisrosei* is found further north and closer to Sacramento, CA (Trock, 2006). Second, there is predicted ancestral introgression from *P. cana* (Hook.) W.A.Weber & Á.Löve into *P. multilobata* (Torr. & A.Gray) W.A.Weber & Á.Löve, potentially explaining why these widespread species place close to each other phylogenetically (Fig. 3; Moore-Pollard and Mandel, 2023). One important note is that *P. cana* and *P. multilobata* have low support in this phylogeny, which may influence our results since the taxon may be misplaced in this clade.

#### Arctic/Alpine

The Arctic/Alpine clade did not experience as much discordance as other clades in the tree. Only one instance of ancestral reticulation is found between the lineages that gave rise to *P. flettii* (Wiegand) W.A.Weber & Á.Löve and *P. contermina* (Greenm.) J.F.Bain, influencing the formation of *P. greenei* (A.Gray) W.A.Weber & Á.Löve (Fig. 7). *Packera greenei* is a species endemic to northern California with deep orange ray florets, a defining character to the species, differing from *P. flettii* and *P. contermina* which have the typical yellow rays seen in most other *Packera* species. These latter species are found in the Pacific Northwest US and Canada and do not geographically overlap with *P. greenei*, so this reticulation event is unexpected though a connection may be found with further investigation.

#### Rocky

There is only one reticulation event in the Rocky Mountain taxa which potentially indicates an ancient hybrid speciation event between *Packera streptanthifolia* var. *rubicaulis* (Greene) Dorn and *P. werneriifolia* (A.Gray) W.A.Weber & Á.Löve ex Trock, producing *P. mancosana* Yeatts, B.Schneid. & Al Schneid. (Fig. 7). *Packera mancosana* is a newly described species that is similar to *P. werneriifolia* in appearance and ecology but differs in general habitat and soil type (Yeatts et al., 2011). Some botanists observing *P. mancosana* in the field noted that it was likely a variable *P. werneriifolia*; however, our study and previous work on the group show that both *P. werneriifolia* and *P. mancosana* do phylogenomically resolve separately from each other, although in the same smaller clade so there may be some species separation (Figs. 2 – 4; Moore-Pollard and Mandel, 2023). Further research delimiting *P. werneriifolia* and *P. mancosana* is needed to support these suspicions.

We note that network analyses, such as PhyloNetworks, are highly sensitive to sampling bias. Even though we sampled 85 out of a targeted 93 *Packera* taxa (∼91% coverage of the genus), missing taxa could influence and change the results. Including the remaining, unsampled *Packera* taxa in the future could help increase our understanding of these relationships. Additionally, limiting the analyses to specified clades can ignore other potential reticulation events occurring across groups. The authors recognize this limit; however, computational requirements of current reticulation network analyses limit the ability to sample more taxa or loci (see Hejase and Liu, 2016). With the recent advances in phylogenetics, we anticipate that tools will soon be developed that are not as computationally intense and will be more efficient with large datasets, so researchers can test for reticulations across entire phylogenies.

Another aspect to consider with hybridization networks is the idea of ghost introgressions, which are ancient introgression events that leave genetic signatures of extinct species in present-day species (Ottenburghs, 2020). Some methods can detect ghost introgressions such as PhyloNetworks (Solís-Lemus and Ané, 2016; Solís-Lemus et al., 2017), but these methods do not specify that the hybrid edge is a ghost introgression, it is up to the researcher to recognize its presence, which can be difficult especially without background knowledge of the group. Even so, accounting for ghost introgressions has important implications for macroevolutionary studies (Ottenburghs, 2020).

## CONCLUSION

We have presented a model for investigating the causes and consequences of nuclear discordance, using an ensemble of approaches to handle paralogs. We aimed to understand how evolutionary processes of ILS and gene flow influence the phylogenetic patterns seen in *Packera*. We found that addressing polyploidy, and ultimately paralogs, increased nodal concordance and enabled us to gain a better understanding of evolutionary relationships in this complex genus. In *Packera*, the tree topology and discordance results indicated that the paralog selection method ASTRAL-Pro outperformed ASTRAL-III and TEO methods. Nonetheless, this trend may not be consistent when tested on other complex plant groups, so we suggest testing multiple paralog selection/pruning methods on one’s dataset to determine the impact of paralogs on your tree topologies and support values. Even after accounting for paralogy, there was an overwhelming amount of discordance present within *Packera*, indicating other biological processes may be influencing our results. In response, we investigated hybridization networks at highly discordant nodes within *Packera*, revealing substantial evidence for ancient hybridization and reticulate evolution, potentially explaining some of the discordance seen within this group. Our work provides further insight into how underlying biological processes can influence species relationships and levels of discordance within phylogenomic studies. Given this, we recommend that future work further investigates these biological processes further as they can impact phylogenomic tree interpretations.

## Acknowledgements

The authors thank Stephen Smith for his assistance in running *bellerophon* and Matthew D. Pollard for bioinformatic help. We also thank the University of Memphis High-Performance Cluster (HPC) administrators, Eric Spangler and Kristian Skjervold, for their assistance with the HPC and willingness to help.

## Author contributions

E.R.M.P. designed the study, generated and analyzed data, and wrote the manuscript. J.R.M. provided guidance on the project and provided edits to the manuscript.

All authors approved of the final version.

## Data Availability Statement

Raw sequence data are available in the National Center for Biotechnology Information (NCBI) Sequence Read Archive (Bioproject: PRJNA978568).

Additional Supporting Information may be found online in the Supporting Information section at the end of the article.

**Appendix S1.** Voucher specimens of the 108 taxa used in this study. Publication status and authorities assigned by IPNI.

**Appendix S2.** Statistics generated from the HybPiper pipeline.

**Appendix S3.** Locus list indicating the loci containing paralogs that were retained (692ortho-para), pruned loci containing paralogs (118para), and orthologous loci not used as input for the TEO pipeline.

**Appendix S4.** A tanglegram of this study’s new 108 taxa tree (left) compared to the original 106 taxa tree (right) from Moore-Pollard & Mandel (2023). Red dash lines indicate highly different topologies between trees.

**Appendix S5.** Phylogeny of ASTRAL-III tree containing 108 taxa with all LPP values shown at the nodes.

**Appendix S6.** Phylogeny of ASTRAL-Pro tree containing 108 taxa with all LPP values shown at the nodes.

**Appendix S7.** Phylogeny of ASTRAL-III tree generated from the results of TEO methods containing 108 taxa with all LPP values shown at the nodes.

**Appendix S8.** Results of Quartet Sampling on the ASTRAL-III tree with all Quartet Sampling values (QC/QD/QI) indicated at the node.

**Appendix S9.** Results of Quartet Sampling on the ASTRAL-Pro tree with all Quartet Sampling values (QC/QD/QI) indicated at the node.

**Appendix S10.** Results of Quartet Sampling on the ASTRAL-III tree generated from the results of TEO methods with all Quartet Sampling values (QC/QD/QI) indicated at the node.

**Appendix S11.** Log likelihood values and taxon lists on the five PhyloNetworks runs: Backbone, Arctic/Alpine, California/Mexico, Eastern, and Rocky Mountains. The number of reticulations with the lowest log likelihood value is considered the best fit.

## Notes

### Competing Interest Statement

The authors have declared no competing interest.

