## Appendices 4-10 for "From paralogy to hybridization: Investigating causes of underlying phylogenomic discordance using the complex genus *Packera* (Senecioneae; Asteraceae)"

Supplemental Table of Contents


| **Name** | **Brief description** | **Page #(s)** |
|---|---|---|
| Appendix S1 | Vouchers of all 108 taxa | Attached (excel sheet) |
| Appendix S2 | General and HybPiper statistics of 108 taxon tree | Attached (excel sheet) |
| Appendix S3 | Locus statistics of TEO pipeline | Attached (excel sheet) |
| Appendix S4 | Tanglegram of 106 taxon tree published in Moore-Pollard & Mandel (2023) and 108 taxon tree from this study | 1 |
| Appendix S5 | Phylogeny of ASTRAL-III tree with all LPP values | 2 |
| Appendix S6 | Phylogeny of ASTRAL-Pro tree with all LPP values | 3 |
| Appendix S7 | Phylogeny of TEO tree with all LPP values | 4 |
| Appendix S8 | Quartet Sampling results across all nodes of ASTRAL-III tree | 5 |
| Appendix S9 | Quartet Sampling results across all nodes of ASTRAL-Pro tree | 6 |
| Appendix S10 | Quartet Sampling results across all nodes of TEO tree | 7 |
| Appendix S11 | Statistics from PhyloNetworks run | Attached (excel sheet) |


**Appendix S1.** Voucher specimens of the 108 taxa used in this study. Publication status and authorities assigned by IPNI.

**Appendix S2.** Statistics generated from the HybPiper pipeline.

**Appendix S3.** Locus list indicating the loci containing paralogs that were retained (692ortho-para), pruned loci containing paralogs (118para), and orthologous loci not used as input for the TEO pipeline.

**Appendix S11.** Log likelihood values and taxon lists on the five PhyloNetworks runs: Backbone, Arctic/Alpine, California/Mexico, Eastern, and Rocky Mountains. The number of reticulations with the lowest log likelihood value is considered the best fit.

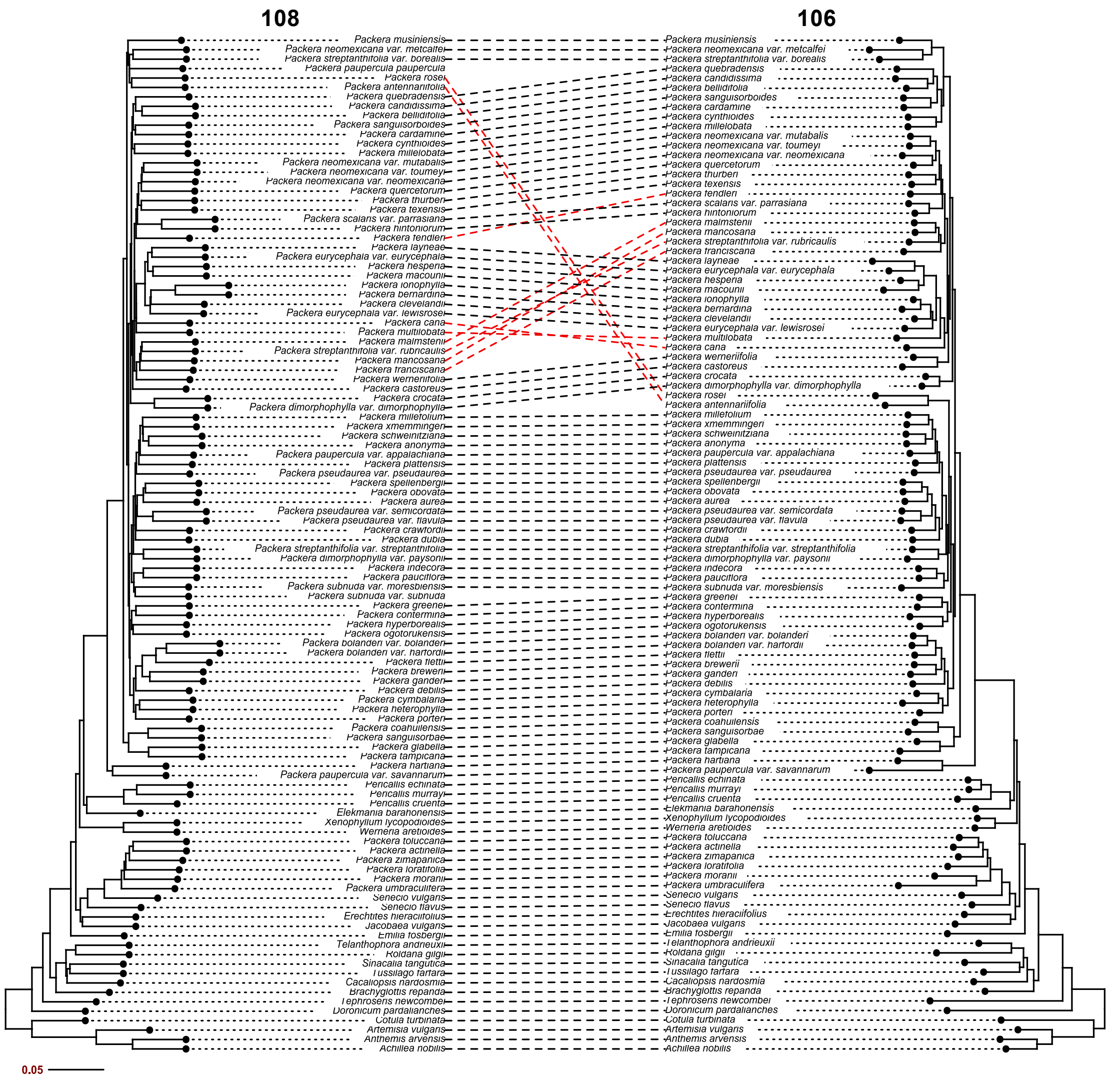


**Appendix S4.** A tanglegram of this study's new 108 taxa tree (left) compared to the original 106 taxa tree (right) from Moore-Pollard & Mandel (2023). Red dash lines indicate highly different topologies between trees.

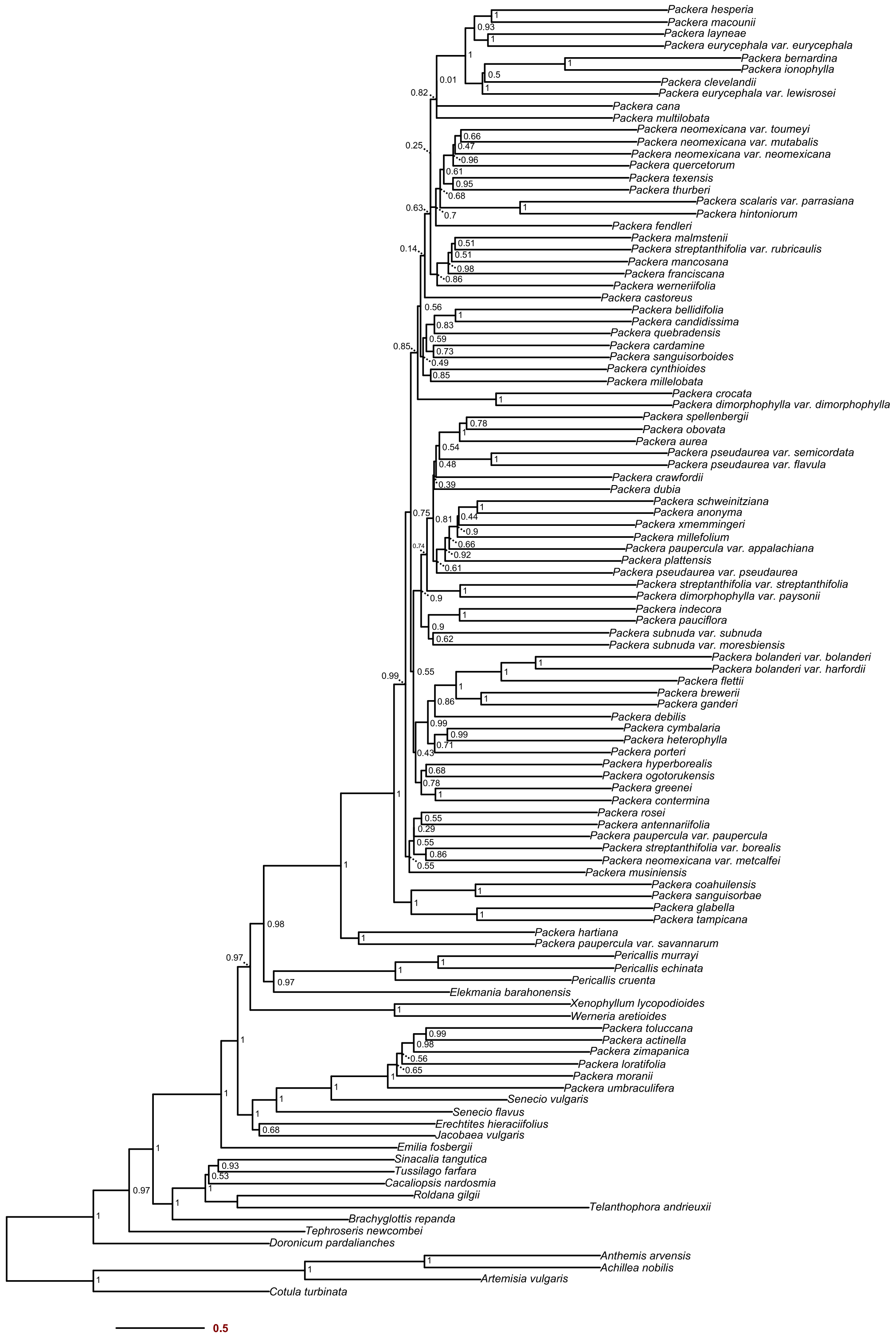


**Appendix S5.** Phylogeny of ASTRAL-III tree containing 108 taxa with all LPP values shown at the nodes.

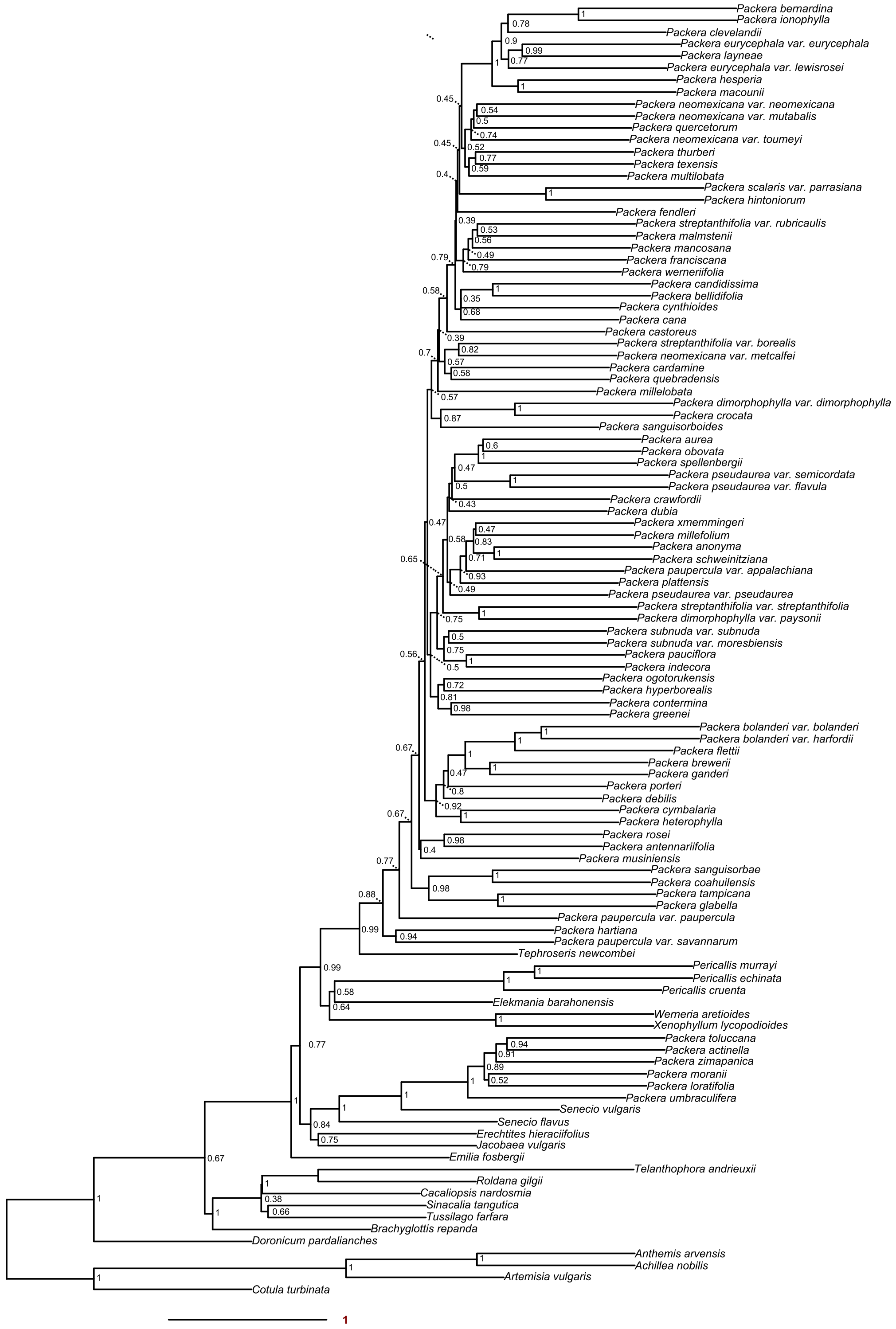


**Appendix S6.** Phylogeny of ASTRAL-Pro tree containing 108 taxa with all LPP values shown at the nodes.

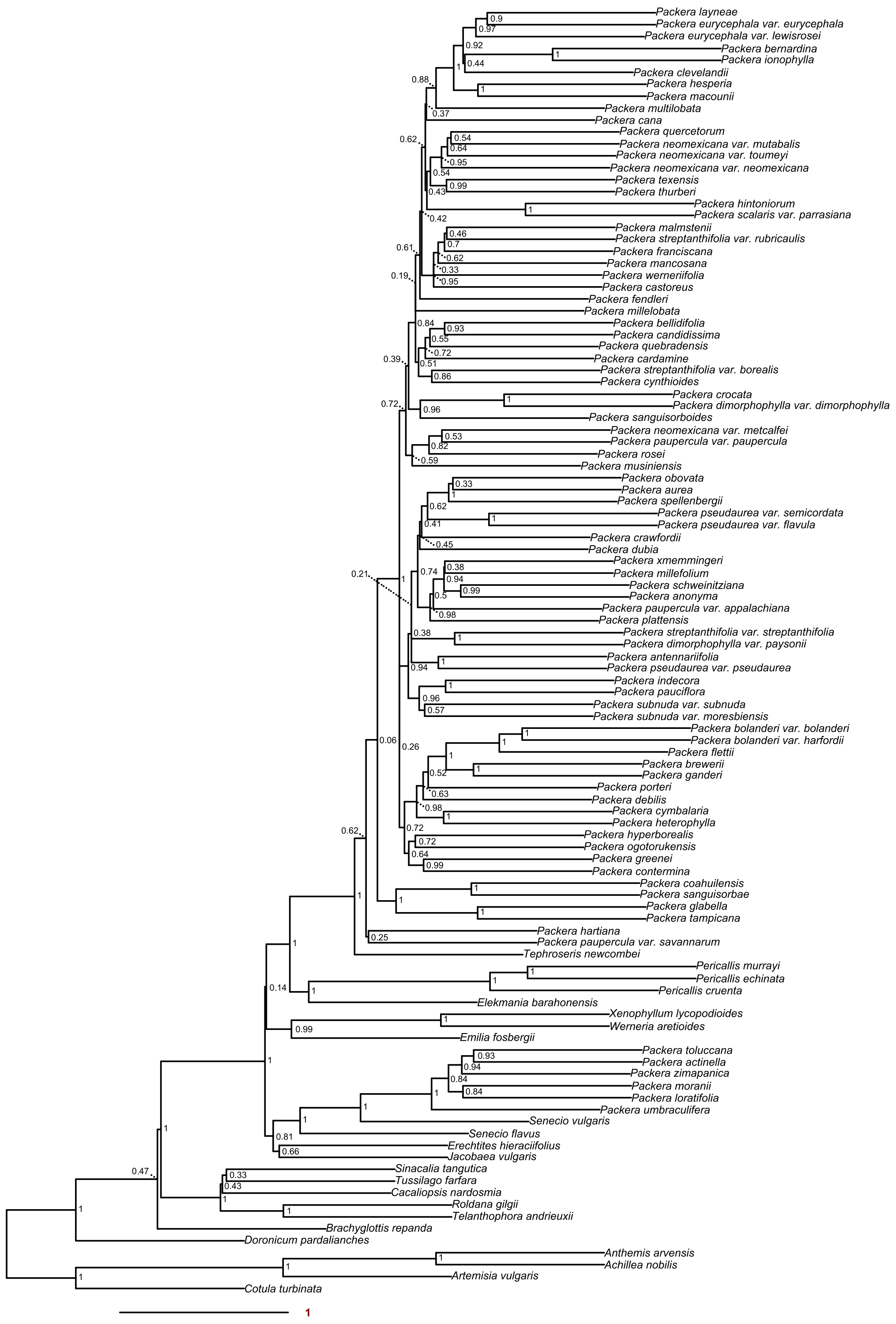


**Appendix S7.** Phylogeny of ASTRAL-III tree generated from the results of TEO methods containing 108 taxa with all LPP values shown at the nodes.

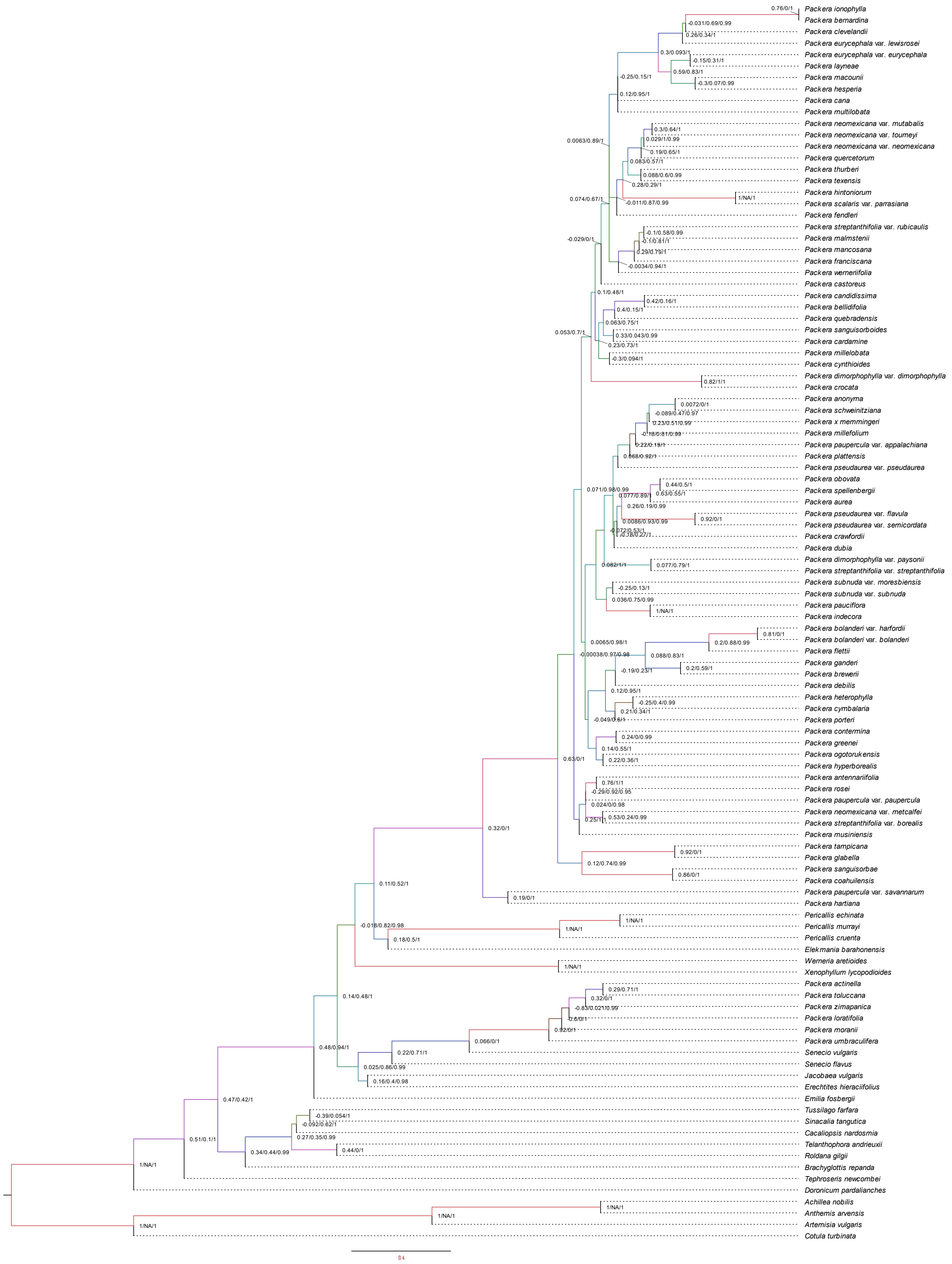


**Appendix S8.** Results of Quartet Sampling on the ASTRAL-III tree with all Quartet Sampling values (QC/QD/QI) indicated at the node.

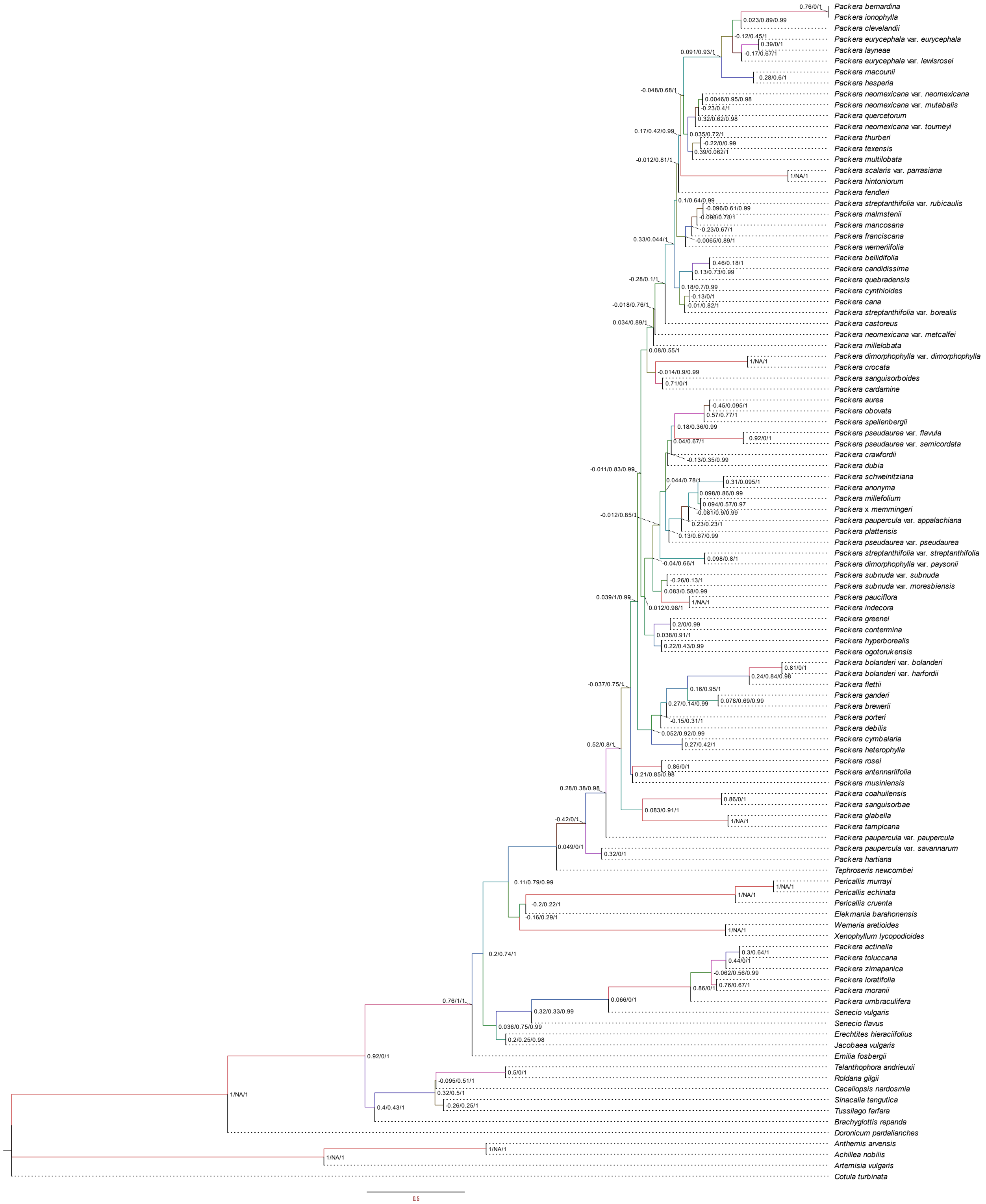

**Appendix S9.** Results of Quartet Sampling on the ASTRAL-Pro tree with all Quartet Sampling values (QC/QD/QI) indicated at the node.

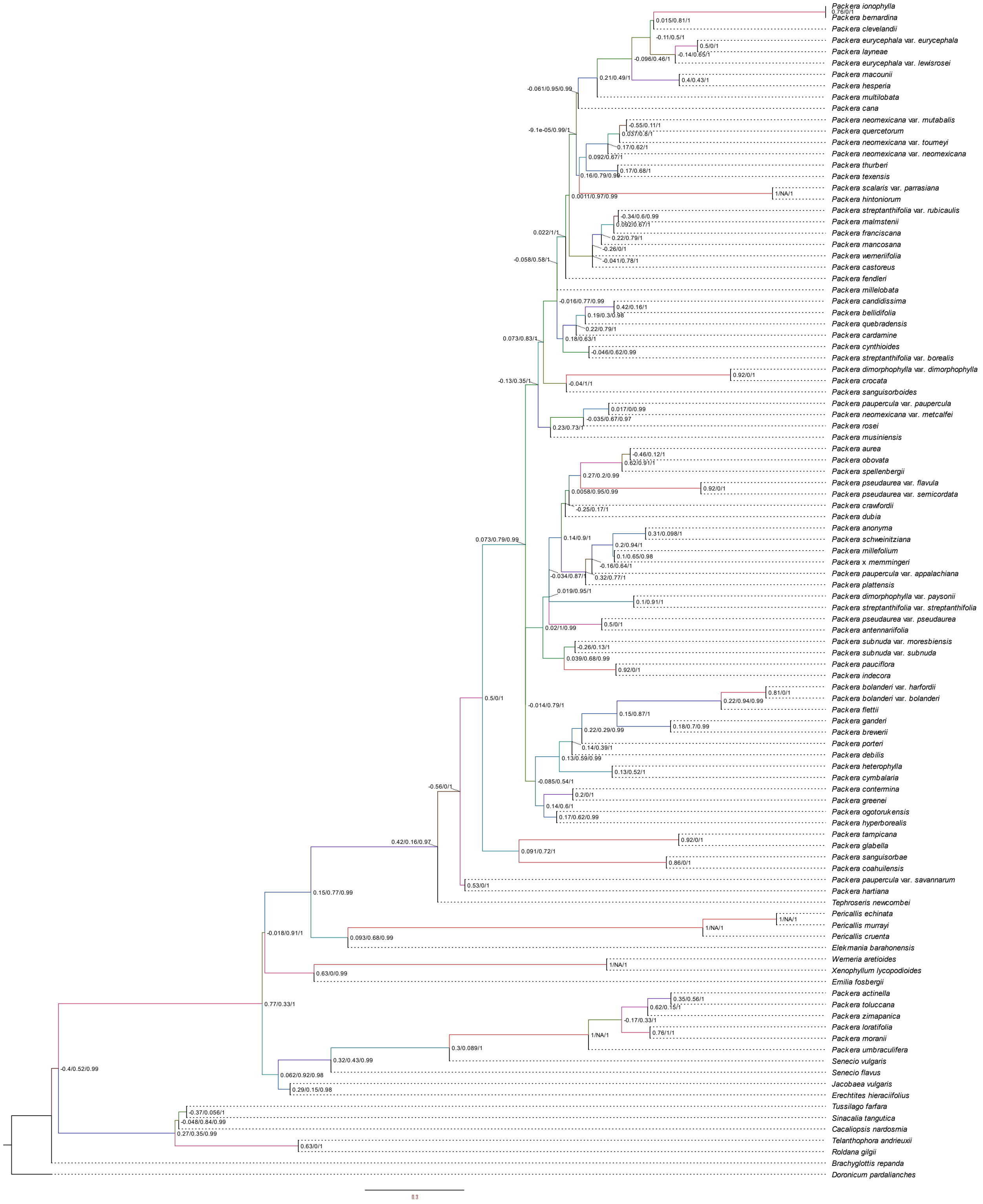


**Appendix S10.** Results of Quartet Sampling on the ASTRAL-III tree generated from the results of TEO methods with all Quartet Sampling values (QC/QD/QI) indicated at the node.
